# The effect of exercise on the protein profile of rat knee joint intra- and extra-articular ligaments

**DOI:** 10.1101/2020.01.09.900142

**Authors:** Yalda A. Kharaz, Helen L. Birch, Alexandra Chester, Eleanor Alchorne, Deborah Simpson, Peter Clegg, Eithne Comerford

## Abstract

Injuries to the intra-articular knee joint ligament (anterior cruciate ligament (ACL)) together with the extra-articular medial collateral ligament (MCL) result in significant joint instability, pain and immobility for the affected individual. Moderate endurance type exercise has been shown to increase ligament strength, however little is known on the effect of short-term high intensity exercise regimes such as treadmill training on the ACL and MCL and whether they may be beneficial to the extracellular matrix (ECM) structure of these ligaments. This study aimed to identify the effect of short-term high intensity exercise on the proteome of the rat ACL and MCL using mass spectrometry. Sprague Dawley male rats (n=12) were split into equal groups of control and exercise animals, which were subjected to high intensity training and followed by proteomic analysis of the ACL and MCL. Knee joint and ligament health was assessed using OARSI scoring or using a validated histological scoring system. Histopathological analyses demonstrated no significant changes in the ACL, MCL or cartilage of the knee joint, indicating that the exercise regime used in this study did not have substantial impact on tissue structure and health of several tissues within the rat knee joint. Some proteins were found to be significantly more abundant in the ACL in the exercised group than the control group. However, no proteins with a significantly different expression were identified between MCL control and MCL exercised groups. The majority of proteins expressed at higher levels in the ACL exercise group were cytoskeletal proteins, ribosomal proteins and enzymes. Several matrisomal proteins were also more abundant such as collagen proteins and proteoglycans in ACL exercise group. In conclusion, our results indicate that short-term high intensity exercise has an impact on ACL ECM protein expression, with the majority of differential expressed proteins being cellular proteins such as actins, ribosomal and heat shock proteins, indicative of metabolic and molecular responses. Further study is necessary to determine the impact of these short-term changes on ligament structure and function.

## INTRODUCTION

Ligaments are short bands of fibrous connective tissues and are responsible for providing a mechanical connection from bone to bone across joints [1]. Their function is to guide and limit normal joint motion, assisted by joint surface geometry and musculotendinous forces [2]. The anterior cruciate ligament (ACL) [3] is located intra-articularly in the mammalian knee joint and is one of the most frequently injured ligaments within this joint. Injuries to the ACL together with the extra-articular medial collateral ligament (MCL), account for over 95% of all multi-ligament injuries in the knee joint [4, 5], resulting in significant joint instability and causing major physical, social [6] and financial implications [7]. ACL injuries can also lead to significant functional impairment in athletes as a result of joint instability and muscle atrophy, and are associated with the development of osteoarthritis [5, 8] leading to a major clinical challenge in orthopaedic medicine [9].

Exercise is known to exert beneficial effects on the musculoskeletal system by enhancing muscle mass, increasing bone strength [10] and contributing to the mechanical strength of ligaments and tendons [11, 12]. In tendons, such as Achilles and patellar tendon, endurance-type exercise has been shown to increase stiffness [13], tensile strength [12, 14] and increases in the cross-sectional area [15-17]. In the mouse Achilles and patellar tendons, it has been demonstrated that short-term treadmill running enhances levels of growth factors such as insulin-like growth factor 1 (IGF-1) [18-20] and transforming-growth factor beta (TGFß) [21, 22]. Together these data, along with increased collagen synthesis observed after both acute exercise and endurance training, suggest that tendon is dynamic in its response to mechanical loading [23-26].

In ligaments, it has been shown that enforced treadmill running has a beneficial effect on the strength of the MCL in rat knees. In the ACL, increased intercellular activity of fibroblasts and decreased average fibril diameters have been observed in adult rats after acute treadmill running, which is indicative of increased collagen metabolism [27]. Endurance treadmill training in rats has been shown to be beneficial for ACL strength and mechanical stiffness[11]. Other studies have shown an increase in extracellular matrix (ECM) components such as elastin microfibrils in ACLs from an exercising dog breed (e.g. greyhound) compared with more sedentary dog breeds (e.g. Labrador retriever) [28].

To date, the comparative response to mechanical loading between the MCL and ACL has not been studied, and its response could be used in potential therapeutic strategies to aid these ligaments with repair and to avoid degeneration. The MCL has been found to heal adequately, whilst the ACL has been found to have poor capacity for healing, even following direct apposition with suture repair [29]. These differences may be due to factors which are mechanical and biologic in origin [29], alterations in the cellular metabolism after injury [30] and to intrinsic cell deficiencies of the ACL [31]. There is a paucity of knowledge on the effect of mechanical loading in terms of exercise between the ACL and MCL and whether it may be beneficial to structural ECM proteins in these ligaments. In this study, we hypothesised that short-term high intensity exercise would result in alterations to the ACL and MCL protein profile which would be structure specific and impact on knee joint health. Therefore, this study aimed to identify the effect of short-term high intensity exercise using treadmill training on the proteome of the rat ACL and MCL. The proteome was assessed using our previously established proteomic workflow with label-free quantification [32]. Knee joint health was evaluated using histology to identify pathological changes to structures within the rat knee joint, including the cruciate and collateral ligaments and articular cartilage.

## METHODS

### Training exercise regime and tissue collection

The study was conducted under Home Office project license number PPL 70/7210. Male Sprague Dawley rats (n=12) were assigned to an exercise (n=6) or control (n=6) group. The rats were eight weeks old at the beginning of the study. All rats were housed in the animal facility for one week prior to the start of the study to acclimatise to their surroundings. The rats were group housed and allowed free cage activity and provided with a standard pelleted rat chow and water ad libitum. The rats in the exercise group were introduced to the treadmill (Linton Instrumentation, UK) over a two-week period before commencing a four-week treadmill-training programme. The training programme consisted of running five days/week on the treadmill at 0° incline with increasing speed up to 17m/s for two 30-minute sessions 30 minutes apart (Supplementary Figure 1). At the end of the four weeks rats were euthanased and the left and right knee joints harvested. The right knee joints were removed and prepared for whole joint histological analysis. The left knee joint was dissected and the MCL and ACL removed for protein extraction and proteomic analysis. The left MCL and ACL samples were snap frozen in liquid nitrogen and stored at −80°C until required.

### Proteomics

ACL and MCL samples were homogenized and proteins were extracted, as previously described, [32] using 4M guanidine-HCl followed by RapiGest™ extraction and the protein concentration of each soluble fraction was measured using the Pierce™ 660 nm protein assay. 30μg of the soluble fraction was subjected to an in-solution digestion on 10 μl Strataclean™ resin (Agilent Genomics, UK) followed by reduction and alkylation in 3 mM dithiothreitol (DTT) and 9 mM iodoacetamide with trypsin at a ratio of 50:1 protein : trypsin [32]. Liquid chromatography tandem-mass spectrometry (LC-MS/MS) analysis was performed using an Ultimate 3000 nano system (Dionex/Thermo Fischer) coupled online to a Q-Exactive Quadrupole-Orbitrap mass spectrometer. 2 μL loading of digests (equivalent to 60 ng peptides) were loaded on to the column on a one-hour gradient with an inter-sample 30 minutes blank as described previously [33, 34].

Proteins were identified using PEAKS studio 8.5 (Bioinformatics Solutions, Waterloo, ON, Canada) using the Uniprot Rat database (UP000002494) as described previously [32, 35]. In brief, instrument configuration was set up as Orbitrap (Orbi-Orbi) and high collisional dissociation (HCD) fragmentation. Parameters used were; 10.0 ppm parent mass error tolerance and 0.01 Da fragment mass error tolerance; trypsin monoisotypic enzyme; one missed cleavage; one nonspecific cleavage; fixed modification, carbamidomethylation; variable modification, methionine oxidation and hydroxylation; and 3 variable PTMs per peptide. Searches were adjusted to confidence score > 50%; protein-10lgP≥ 20, 1% false discovery rate (FDR) and unique peptides ≥ 2. Label free (LF) quantitative analysis was performed firstly by comparing both the ACL and MCL control and exercise groups together. After that, pair-wise comparisons were performed between the samples in the control and exercise groups on Progenesis^QI^ software (Waters, Elstree Hertfordshire, UK) [33, 36]. In brief, the spectra for each feature was searched against the UniProt Rat database on Mascot (v2.6 Matrix Science, London, UK) using the same parameters. Identified peptides hits were re-imported and assigned to proteins and filtered at a threshold score corresponding to a 1% FDR. Identification of protein with two or more peptides were used for quantification, and greater than two-fold abundance with a FDR adjusted p-values <0.05 were considered to be significant. The proteomics data set for this study has been deposited in the ProteomeXchange Consortium via the PRIDE [37] partner repository (identifier PXD016516).

### Gene Ontology, Pathway Enrichment Analysis and Protein Network Analysis

Further analysis of the proteomic data was only performed on ACL control and exercise groups as no significant changes were found between MCL control and exercise group.

Significant abundant proteins quantified using proteomic analysis between ACL control and exercise groups were used to assess the gene ontology (GO), pathway enrichment analysis and protein network analysis. Interactions between proteins were visualised using bioinformatics software to gain a broader understanding of functional changes derived from the ACL exercise group. ToppGene software was used for functional enrichment analysis of the up-regulated proteins in the ACL exercise group with Benjamini-Hochberg false discovery rate adjusted *p*-value cut-off < 0.05 [38]. Biological process GO terms and associated FDR values generated through ToppGene were then summarized and formed into a network of processes using REViGO [39], which was then exported to Cytoscape software [40] to generate interactive graphs. In addition, Strings Bioinformatics tool [41] was used for GO and to produce a further network to visualise connections between up-regulated and down-regulated proteins between ACL control versus exercise group, as described previously [35].

Using the same significant data sets, the canonical pathways, further networks and biologic process pathways were produced using Ingenuity Pathway Analysis software (IPA, Ingenuity Systems, Redwood City, CA, USA). The accession and *p*-values of those significant proteins were used to form a core analysis as described previously [35].

### Western Blotting Validation of Beta-actin Abundance

Western blotting was performed to validate the up-regulation of beta-actin abundance between ACL control and exercise groups using previously established methods [42]. In brief, 10 μg of ACL control and exercise samples were electrophoresed and separated on a pre-cast 12 well gel (Bio-Rad Criterion 10% TGX). Separated proteins were then transferred to a nitrocellulose membrane and blocked with 10 ml LICOR Odyssey^®^ blocking buffer (LI-COR, Cambridge, UK) for 1 hour at room temperature. Subsequently primary antibody (β-actin, Abcam, ab8227, Cambridge, UK) was added to the membrane at a 1:1000 dilution directly into the buffer and allowed to incubate overnight at 4°C. The membrane was washed and incubated in a secondary goat anti-rabbit (LI-COR, IRDye® 680RD Goat anti-Rabbit IgG, Cambridge, UK) at 1:20,000 dilution was incubated for one hour at room temperature. The membrane was imaged by Odyssey^®^ LI-COR CLx imaging system at a wavelength of 700 nm. As a normalising control, GAPDH was also probed following the same steps as above using GAPDH primary antibody (Sigma, Poole, UK) at a 1:1000 dilution with goat – anti rabbit secondary at 1:20,000 (LI-COR, IRDye® 800CW Goat anti-Rabbit IgG, Cambridge, UK). ImageJ software (http://rsbweb.nih.gov/ij/) was used to quantify bands using densitometry. Results were normalised to GAPDH loading control as reported previously [43].

### Histopathological examination of the knee joints

Surrounding soft tissues were removed from the right rat knee joints leaving only the joint capsule and its contents. The dissected joints were stored in 4% paraformaldehyde for 24 hours. Samples were then decalcified for eight weeks in a solution of 25 g EDTA in 175 mls distilled water (pH= 4-4.5) [44] and embedded coronally in paraffin wax and cut at a thickness of 6 μm using a microtome (HM355S, Thermofischer). Sections from the paraffin blocks that were cut prior to the femoral condyles becoming visible were discarded and thereafter three cross-sectional cuttings were mounted per slide. Cutting of the sample paraffin blocks continued throughout the joint until the femur began to disappear visually from samples [45]. Collected sections were stained with haematoxylin & eosin (H&E) [46] and toluidine blue & fast green [47] and histologically scored for the knee joint and knee collateral and cruciate ligament. Sections were scored by two observers blind sighted to the samples using a validated OARSI grading system for rat knee joints [48]. In brief, grade 0= normal, grade 1= minimal degeneration of articular cartilage 5-10% affected, grade 2= mild degeneration, 11-25% affect, grade 3= moderate degeneration, 26-50%, grade 4= marked degeneration, 51-75% affected, grade 5= severe degeneration, 76-100% affected. Medial and lateral aspects of the tibia and femur were scored individually across the whole joint compartment producing a maximum (most severe) grade and overall “average” maximum grade in each group of rats. In addition, a mean score was produced for each joint and these similarly were used to produce an overall ‘average’ mean grade in each group of rats [49]. The scoring for ligaments was adapted from Kharaz, Canty-Laird [50] and scoring was performed based on strength of ECM staining, cell hypertrophy, cell clustering, loss of alignment and ossification and were graded from 0-4 based on the extent of changes ((0= 0% increase; normal, 1= 5-25% increase; mild abnormality, 2=26-50% increase; moderate abnormality, 3= 51-75% increase; marked abnormality, 4= 76-100% increase; severe abnormality). Inter and intra-observer variability was calculated with Cohen’s Kappa statistics using an online software tool: (http://www.statstodo.com/CohenFleissKappa_Pgm.php).

### Statistical analysis

Statistical analysis for proteomic label free datasets was performed by Progenesis^QI^ on all detected features using transformed normalised abundances for one-way ANOVA as previously described [32, 35]. Identification of proteins with two or more peptides, greater than two-fold abundance and with a *q* value (*p*-value adjusted to FDR) <0.05 were considered significant. Quantitative analysis was initially performed by comparing the four groups of samples together. After that, pair-wise comparison were performed between ACL control and ACL exercise, MCL control and MCL exercise, ACL control and MCL control and ACL exercise and MCL exercise as has been used with similar data previously [35]. Normal distribution for the histological and Western blot data sets was assessed with GraphPad Prism (Version 7, GraphPad Software, USA) using a Kolmogorov-Smirnov test. A one-way ANOVA with Bonferroni *post-hoc* test was performed on histological scoring results between the cruciate and collateral ligaments and t-test was performed on mean and maximum OARSI joint scores and Western blot analysis.

## RESULTS

The bodyweights of the rats at the start of the study did not differ significantly between groups (control – 432.8 ± 18.9 g; exercise −413.7 ± 11.7 g). Both groups of rats gained weight during the study (control – 85.0 ± 32.6 g; exercise - 72.7 ± 17.1 g) but this was not significantly different between groups and the bodyweight of the rats at the end of the study (control – 517.8 ± 34.8 g; exercise – 486.3 ± 22.4 g) were not significantly different between groups. Inclination to run on the treadmill varied between rats. The better runners reached a top speed of 17m/s while the poorest runner reached a top speed of 10m/s. This difference resulted in a range of total distance covered by individual rats in the exercise group from 10058.9 to 17506.3 metres.

### Histological findings

In general, minor changes were observed histologically in knee joint health between the control and exercise groups. Histological observation of staining of the cruciate ligaments from control and exercise groups showed a normal alignment with variation in fibre orientation and a similar intensity of toluidine blue staining (Figure 1A, 1Aa, D and Da). For the collateral ligaments, the level of ECM staining between the control and the exercise group were similar with no obvious disorganisation of fibres alignment (Figure 1B, 1Ba, E and Ea). Histological staining of the articular cartilage knee joints showed a smooth undisrupted articular cartilage surface with none to minor degradation and lesions observed in both control and exercise group (Figure 1C, Ca, F and Fa). However, in the right joint from one rat from the exercise group, which ran at lowest a distance of 10058.9 metres, lesions were observed on the surface of the articular cartilage as highlighted in Figure 1F and 1Fa. Histological analysis resulted in average scoring in the ACLs of 3.53 ± 0.92 and 3.66 ± 1.55 for the control and exercise groups respectively. For the MCLs, scores reached 1.69 ± 1.02 in the control group and 2.54 ± 1.03 in the exercise group (Figure 4G). No significant difference was found between control and exercise groups in either cruciate and collateral ligaments (*p*=0.08, *p*=0.19).

**Figure 1.**
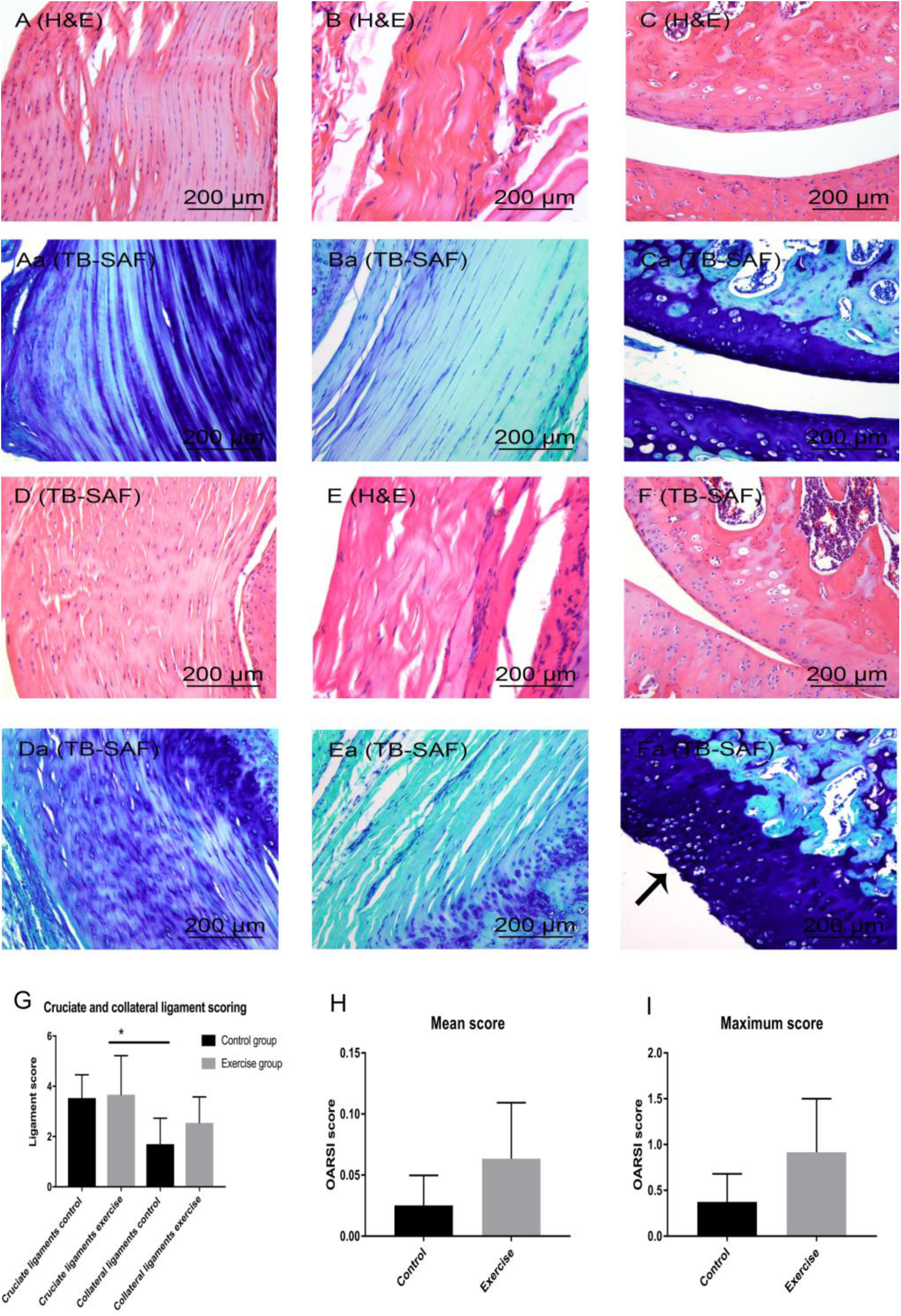
Histological comparison between ACL (A, Aa), MCL (B, Ba), cartilage (C, Ca) control groups and ACL (D, Da), MCL (E, Ea) and cartilage exercise (F, Fa) groups. No significant difference was found in the ligament score (G) and OARSI mean (H) and maximum (I) score between the control and exercise groups.

**Figure 2.**
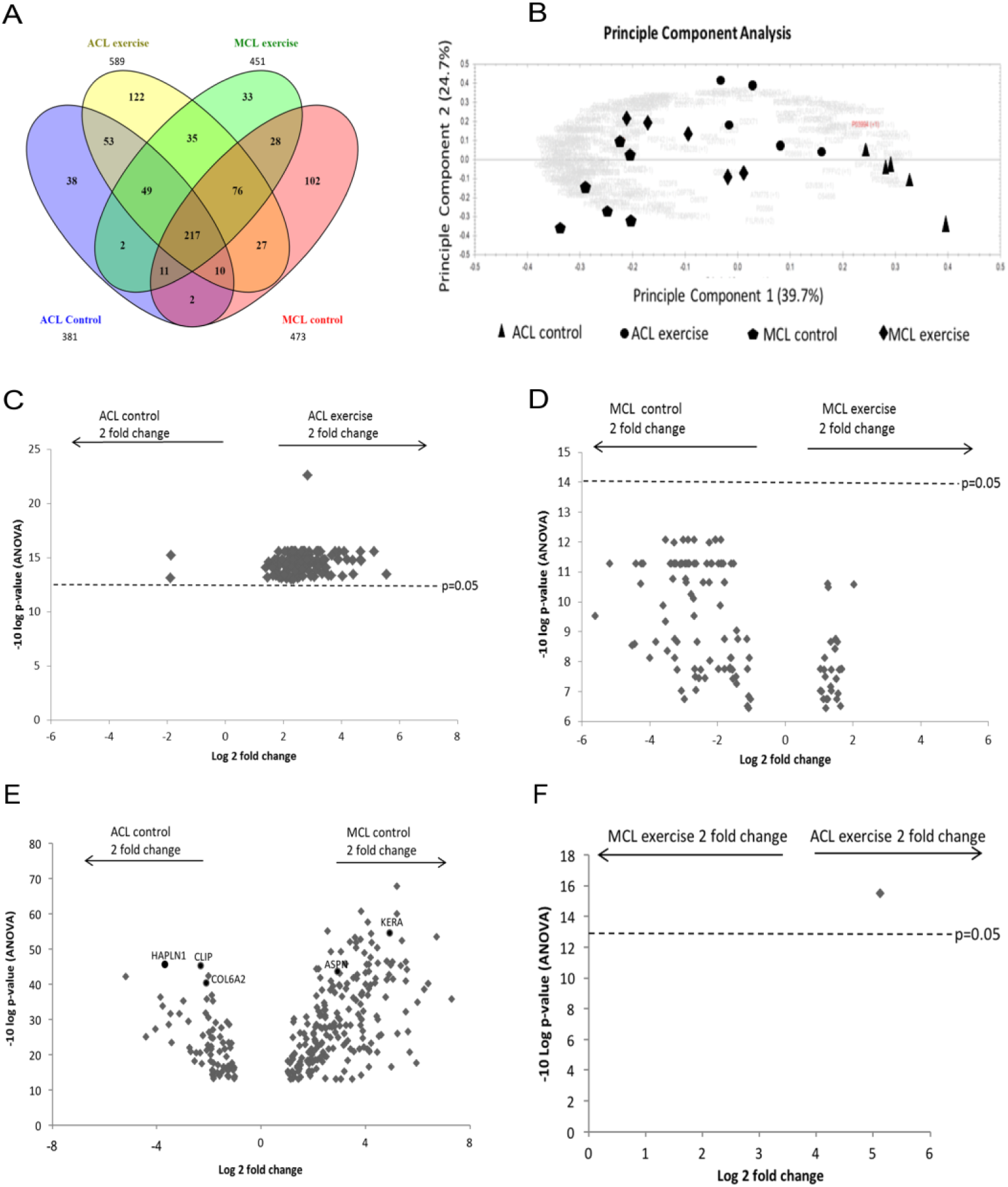
Qualitative and quantitative differences in proteins identified between ACL, MCL control and exercise groups. (A) Venn diagram demonstrating the total number of proteins identified following MS in both ACL and MCL control and exercise group as well as common proteins between the groups. (B) Principle component analysis between ACL and MCL control and exercise groups produced by Progenesis^QI^ after ANOVA analysis with identified proteins at p-value < 0.05. (C-F) volcano plots (−10lgP of FDR adjusted p-value vs log2 fold change). (C) ACL control vs ACL exercise, (D) MCL control vs exercise, (E) ACL control vs MCL control and (F) MCL exercise vs ACL exercise. Volcano plots of quantified proteins in C, D, E indicated up-regulation and down-regulation of proteins with up-fold and down-fold change with significance. This was not the case in volcano plot D as quantified proteins had a p-value (adjusted to FDR)> 0.05.

**Figure 3.**
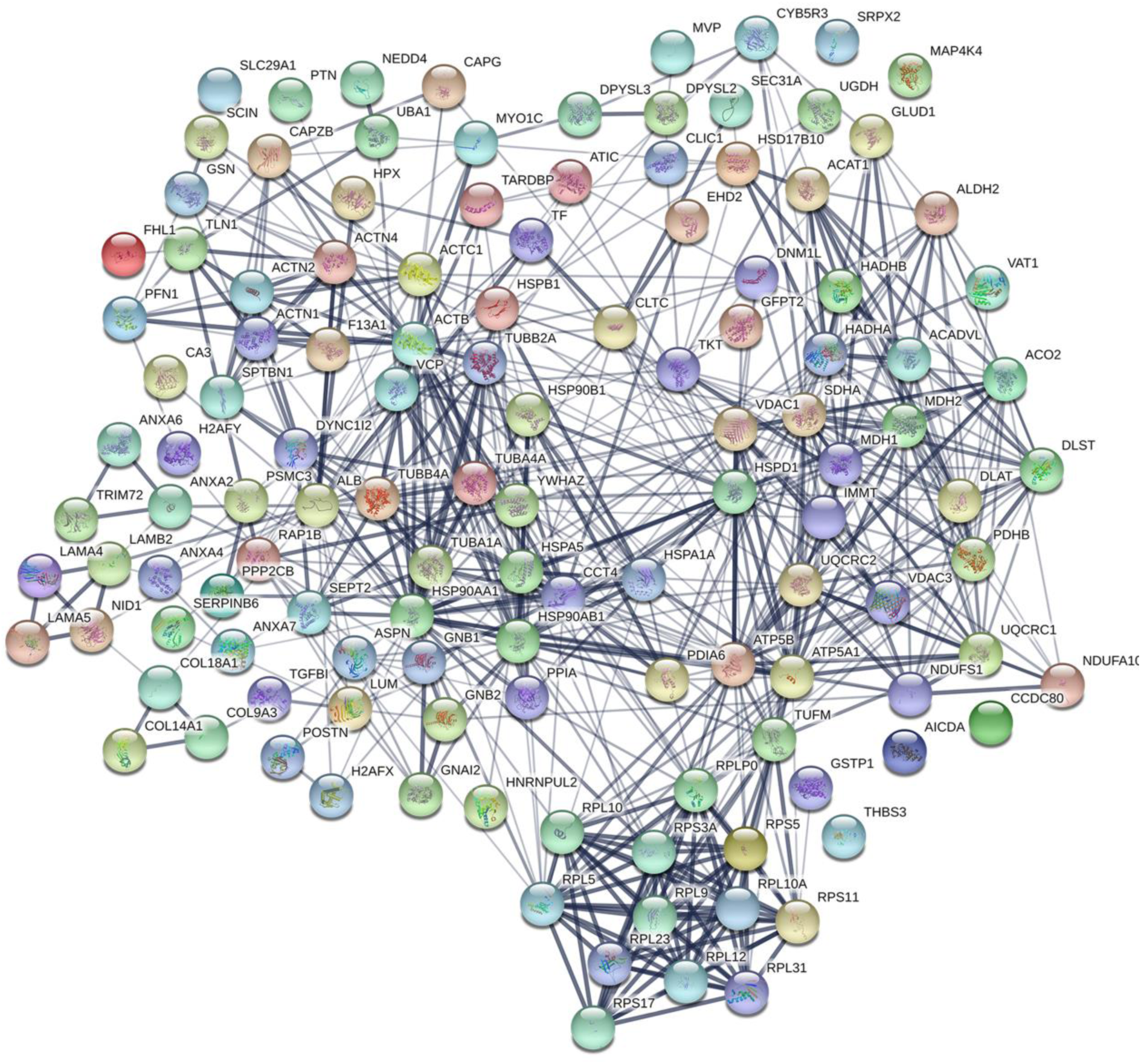
String analysis of upregulated proteins in ACL exercise group versus ACL control group. The above figure shows the greatest linkage of proteins predominantly involves those associate with ribosomes, also there is further linkage of actins, heat shock proteins and collagens. The main principal gene ontology processes were identified as metabolic (*p*= 7.08e-16) and cellular (*p*=2.05e-11).

**Figure 4.**
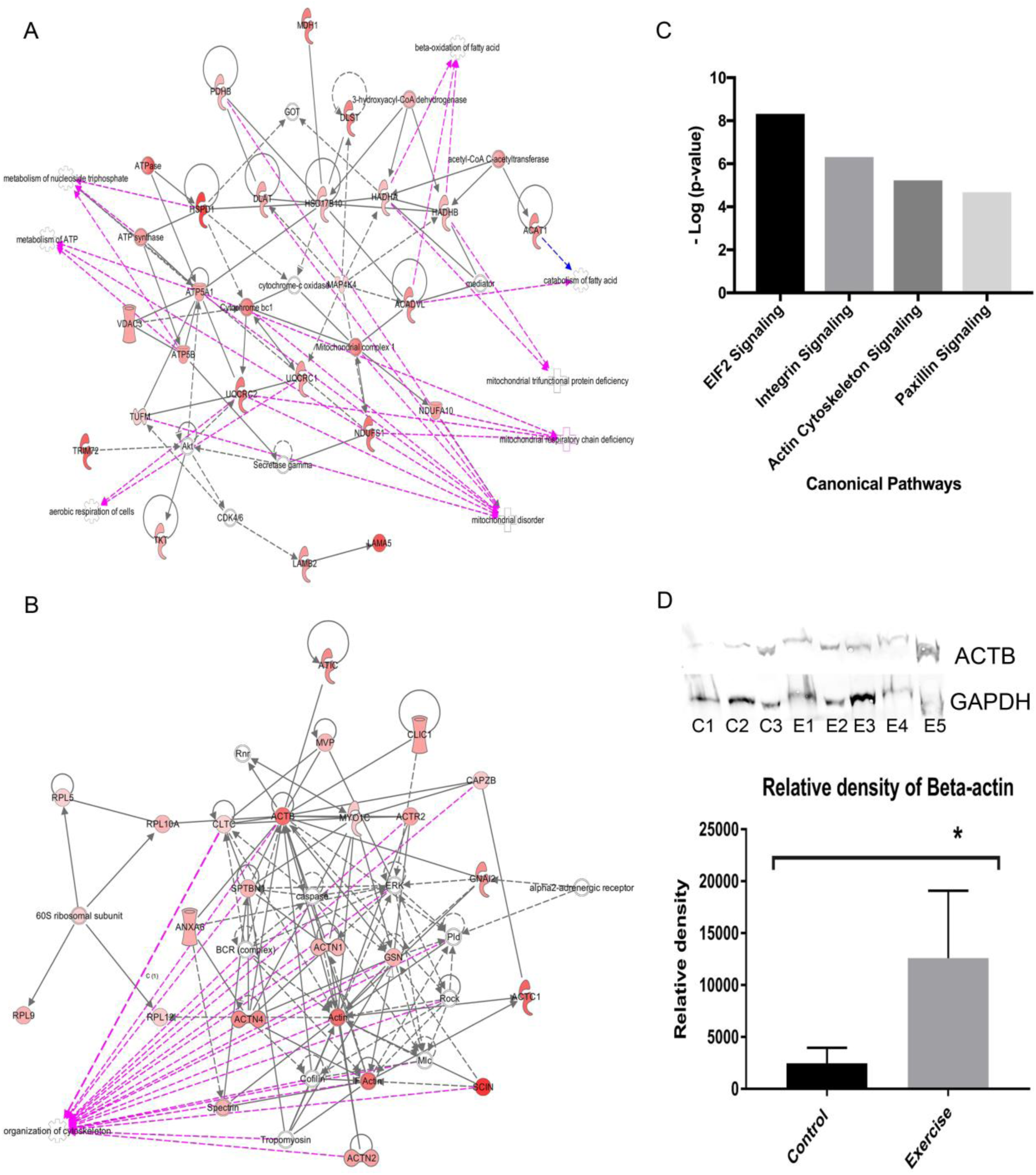
IPA analysis between differential abundant protein between ACL control and exercise group significant networks were related to metabolic, disease development (**A**), and cell signalling, posttranslational modification and protein synthesis (**B**). Red nodes, greater protein abundance in the ACL exercise group; white nodes, proteins not differentially abundant between the ACL exercise and control group. Intensity of colour is related to higher fold-change. Key to the main features in the networks is shown. Significant functions related to network 1 included metabolism of ATP, metabolism of nucleoside triphosphate, beta-oxidation and catabolism of fatty acid, mitochondrial disorder respiratory chain and trifunctional protein deficiencies (**A**, *p* < 0.0001). Diseases and functions related to network 2 included organisation of the cytoskeleton (**B**, *p* < 0.0001). (C) A number of canonical pathways shown to be upregulated in the ACL exercise compared to the ACL control group. (D) Western blot analysis between of Beta-actin in ACL control and ACL exercise. Statistical differences were assessed between the ACL control and exercise group using a T-tests.

An average OARSI score of 0.37 ± 0.3 and 0.92 ± 0.54 was achieved for the control and exercise group, respectively. Overall, the mean OARSI scores for the control and exercise group were calculated at 0.025 ± 0.024 and 0.063 ± 0.045 respectively. This difference between the two groups was not statistically significant (*p*=0.072) (Figure 4H and 4I).

### Proteomics

#### 1) Protein concentration and identification

The average protein content (μg/ mg wet wt) of 24.6, 23.1, 30.1 and 23.1 measured for ACL and MCL control and ACL and MCL exercise groups, respectively was not significantly different (Supplementary Figure 2). A total number of peptides of 4065, 5135, 5517, 4546 assigned to 381, 473, 589 and 451 proteins were identified in ACL and MCL control and ACL and MCL exercise, respectively (Figure 2A, Supplementary Table 1). A higher number of unique and total proteins were identified in ACL exercise group in comparison to ACL control group, however in the MCL a similar number of total and unique proteins were identified in both control and exercise group (Figure 2A).

#### 2) Quantitative label-free (LF) analysis

Quantitative LF analysis demonstrated a set of 332 proteins within the four groups with a fold change >2 and unique peptides >2 (Supplementary Table 2). Principle component analysis (PCA) was used to identify the major variance between the groups. This analysis revealed that the ACL and MCL control samples were distinctly grouped, whereas ACL and MCL exercise samples were clustered closer together (Figure 2B).

Quantitative differences between ACL control and exercise group samples resulted in 124 proteins that were significantly different. Of these proteins, 122 were more abundant in the ACL exercise group and two proteins were more abundant in ACL control group (Figure 2C, and Supplementary Table 3). The majority of significantly more abundant proteins in ACL exercise groups were cytoskeletal, ribosomal and enzymes (Table 1). Several abundant matrisomal proteins such as collagen alpha-3 (IX) chain, collagen type XVIII alpha 1 chain, collagen alpha-1(XIV) chain, asporin, lumican, thrombospondin-3, periostin and TGFß were found to be up-regulated in ACL exercise group. A summary of the classification of these proteins is provided in Table 1.

**Table 1.** Classification of differentially abundant proteins (identified using ≥ 2 peptides, >2-fold change, FDR adjusted *p*<0.05) in the ACL exercise group compared ACL control group.

|  |  | **Higher in**<br>**ACL exercise**<br>**group** | **Higher in**<br>**ACL**<br>**control**<br>**group** |
| --- | --- | --- | --- |
| ECM Proteins | Collagens | 3 | 0 |
|  | Glycoproteins | 6 (4.8%) | 0 |
|  | Proteoglycans | 2 (1.6%) | 0 |
|  | ECM affiliated proteins | 5 (4%) | 0 |
|  | **Total number of proteins** | **16 (12.9%)** | **0** |
| Non-ECM<br>proteins | Cytoskeletal | 17 (13.7%) |  |
|  | Histones | 2 (1.6%) |  |
|  | Cell membrane | 7 (5.6%) |  |
|  | Ribosomal | 13 (10.5%) |  |
|  | Enzymes | 41 (33.1%) |  |
|  | Transcription/translation | 3 (2.4%) |  |
|  | Other/unknown | 23 (18.5%) | 2 |
|  | **Total number of proteins** | **106 (85.5%)** | **2 (1.6%)** |

No statistically significant differences in proteins abundance were identified between MCL control group when compared to the MCL exercise group (Figure 2D and Supplementary Table 4) as all proteins had a FDR adjusted *p*-values greater than 0.05.

When the ACL control was compared to MCL control group samples, 73 proteins were more abundant in ACL control and 217 in MCL control (Figure 2E and Supplementary Table 5). The ACL control group samples were more abundant in fibrocartilaginous proteins such as cartilage intermediate layer protein and hyaluronan and proteoglycan link protein 1, whilst the MCL control group samples had more asporin and keratocan (Figure 1E). Between the ACL and MCL exercise groups only HAPLN was found to be significantly upregulated in the ACL exercise group (Figure 2F and Supplementary Table 6).

### Gene Ontology and Ingenuity Pathway Analysis

Following Cytoscape software analysis of the significantly upregulated proteins in the ACL exercise group, they were found to be proteins mostly associated with respiration and metabolism (Supplementary Figure 3). In addition, gene response to stimuli, protein localisation and cell migration were also significantly upregulated. String analysis demonstrated some similarities with the most predominant linkage involving the ribosomal proteins in ACL exercise group (Figure 3). Further linkage was also seen between the heat shock proteins, actins and collagens. The most common biological processes highlighted by the String analysis software included metabolic (*p*= 7.08e-16) and cellular (*p*=2.05e-11) pathways (Figure 3).

The IPA of the differentially abundant proteins in ACL exercise group compared to ACL control group generated several networks that were enriched (Figure 4A and 4B). According to the top scoring networks, the differentially expressed proteins were associated with metabolic and disease development, cell signalling and post-translational modifications (Figure 4A). Proteins that were found to be enriched included metabolism of ATP and nucleoside triphosphate, aerobic respiration, mitochondrial disorder, respiratory chain and trifunction protein deficiency and organisation of cytoskeleton (Figure 4B). Significant IPA canonical pathways that were upregulated included eukaryotic initiation factor, integrin and actin cytoskeletal and paxicillin signalling (Figure 4C).

Western blot analysis of beta-actin abundance was in agreement with the mass spectrometry data and was significantly greater (*p* = 0.017, (Figure 4D) in the ACL exercise group than ACL control group.

## DISCUSSION

This is the first study to measure the effect of an imposed and controlled exercise regime on the proteome of the rat intra-articular ACL and extra-articular MCL. Our findings demonstrate that short-term (4 weeks) treadmill training influences intra-articular ACL protein expression, but not that of the extra-articular MCL compared to control groups. These changes in protein expression in the ACL as a response to exercise may contribute to a protective or degenerative role in these ligaments. The health of the knee joint, as assessed by histopathological examination, demonstrated no significant differences in the ACL, MCL and cartilage in the exercise groups compared to rats undertaking only cage activity suggesting that the exercise is not detrimental to the soft tissues of the joint. Additional studies are now required to assess biomechanical alterations of the ligament with exercise.

For our proteomic analysis, we used label-free quantification to identify differentially abundant proteins between control and exercise group of both ligaments and between ACL and MCL tissues. This produced 124 significant proteins that were more abundant in ACL exercise than ACL control group. However, no statistically significant differential proteins were identified between MCL control and exercise group. The differences found in protein expression in this study between ACL and MCL exercise groups may be due to altered mechanical loading between the intra- and extra-articular ligaments. In humans, during athletic tasks such as jump landing, the ACL has been found to exhibit greater loading and strain and greater contribution to knee restraint, in comparison to the MCL [51]. In the current study, the rat MCL may be similar to the human MCL and may be subject to less strain compared to the ACL. The exercise regime was given in a straight line, with no twisting or turning, which may limit the load on the MCL and as consequence resulted in different protein expression to that seen in the ACL during exercise. Furthermore the intra-articular ACL is also exposed to cytokines and other mediators released from other joint tissues into synovial fluid, which may also have led to an altered protein profile within the ACL with exercise [52]. Further studies are required to understand the rat knee joint loading during *in vivo* tasks and may provide insight that enhances the efficacy of injury prevention protocols.

Our proteomic analysis between ACL control and exercise group samples demonstrated an increase in mainly cellular proteins such as tubulins, ribosomal and heat shock proteins. We also found actins to be abundant in the ACL exercise group, which were then validated through western blot analysis. Actin participates in many cellular processes such as muscle contraction, cell motility, division, cytokinesis and signalling, where many of these processes are mediated by extensive and intimate interactions of actin with cellular membranes [53, 54]. In tendon, the disruption of the actin cytoskeleton has been found to decrease tissue elastic modulus during development [55]. Therefore, the increased actin protein found in our study in ACLs following exercise could contribute to the improved tissue mechanical properties. Whilst the majority of abundant proteins were cellular associated, several matrisomal collagens, proteoglycan and glycoprotein proteins such as collagen type IX, XIV and XVIII, lumican, asporin, periostin, thombrospondin-3 and TGFß were also upregulated in the ACL exercise group. The exact role and mechanism of these matrisomal proteins is not known after exercise, but the presence of collagen type IX may indicate a chondrocytic phenotype of ACL. This corresponds with another study in mouse Achilles tendon where intense exercise resulted in cartilaginous changes [56]. The upregulation of TGFβ found in this study could indicate local release in the ACL tissue and agrees with previous tendon exercise studies where elevations of TGFß have been demonstrated in response to exercise [25, 57]. In tendon, mechanical loading following exercise has been shown to release active TGFß, which has been demonstrated to regulate ECM protein expression such as collagen type I [25], proteoglycans [58], and also microRNA molecules with known roles in cell proliferation and extracellular matrix synthesis [59]. The upregulation of TGFß in the ACL exercise group may be associated with regulation of ECM proteins and is likely to stimulate many anabolic pathways that control exercise-mediate ACL adaption.

Gene ontology revealed that metabolic and cellular processes were overrepresented in the ACL exercise group in comparison to the control group. This was also evident using IPA, where the analysis of differential networks identified significant pathways in relation to metabolic development and cell signalling. Ingenuity pathway analysis (IPA) also showed upregulation of several canonical pathways including eukaryotic initiation factor 2 (EIF2) and integrin signalling. Eukaryotic initiation factor 2 signalling enhances the initiation of translational and transcriptional activators [60]) and integrins play a crucial role in linking the ECM to the cytoskeleton playing a role in mechanotransduction of muscle and tendon [57, 61]. In the current study, the exact role of the signalling factors in the ACL exercise group cannot yet be elucidated and additional studies are required to understand whether induced activation of these pathway aid in the organisation of ACL ECM.

In conclusion, we have shown for the first time the effect of strenuous exercise on intra- and extra-articular knee joint ligaments. This study demonstrated that short-term strenuous treadmill exercise impacts ACL protein expression, whilst MCL proteome is not altered. These differences in response may be due to a difference in mechanical loading and previously identified structural and ECM compositional difference between the two tissue types [50]. Although increases in matrisomal associated proteins were observed between ACL control and exercise group, the majority of differentially expressed abundant proteins were cellular, indicative of an intracellular response. Whether these changes are protective or degenerative in ACL is yet to be elucidated.

## ACKNOWLEDGEMENT

The authors wish to thank the Royal National Orthopaedic Hospital Charity, Stanmore for funding the study and the University of Liverpool Technology Directorate for proteomic funding.

## DECLARATION

The authors have declared no conflict of interest.

