## Supplementary Figure 1 for "The effect of exercise on the protein profile of rat knee joint intra- and extra-articular ligaments"

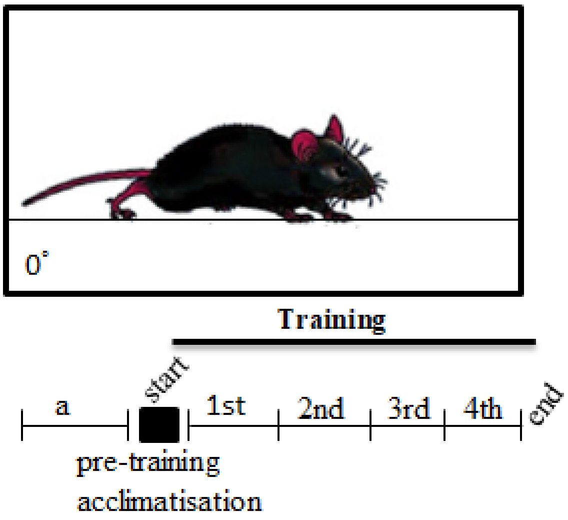


Supplementary Figure 1. Experimental set up of endurance training regime. Rats ran on flat treadmill and were acclimatised 2 weeks prior to training, which was followed by 5 days per week, for total of 1 hour per day, for four consecutive weeks at a speed with a range of speed between 10m/min and 17m/min 14m/min.

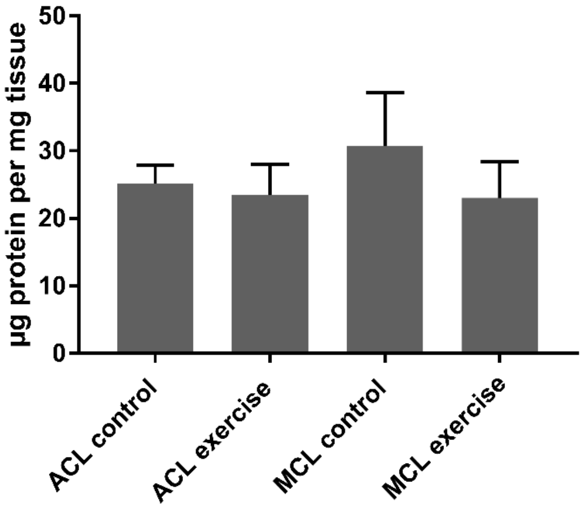


Supplementary Figure 2. (A) Protein concentration yielded between control and exercise groups. Values are mean and error bars represent SD.

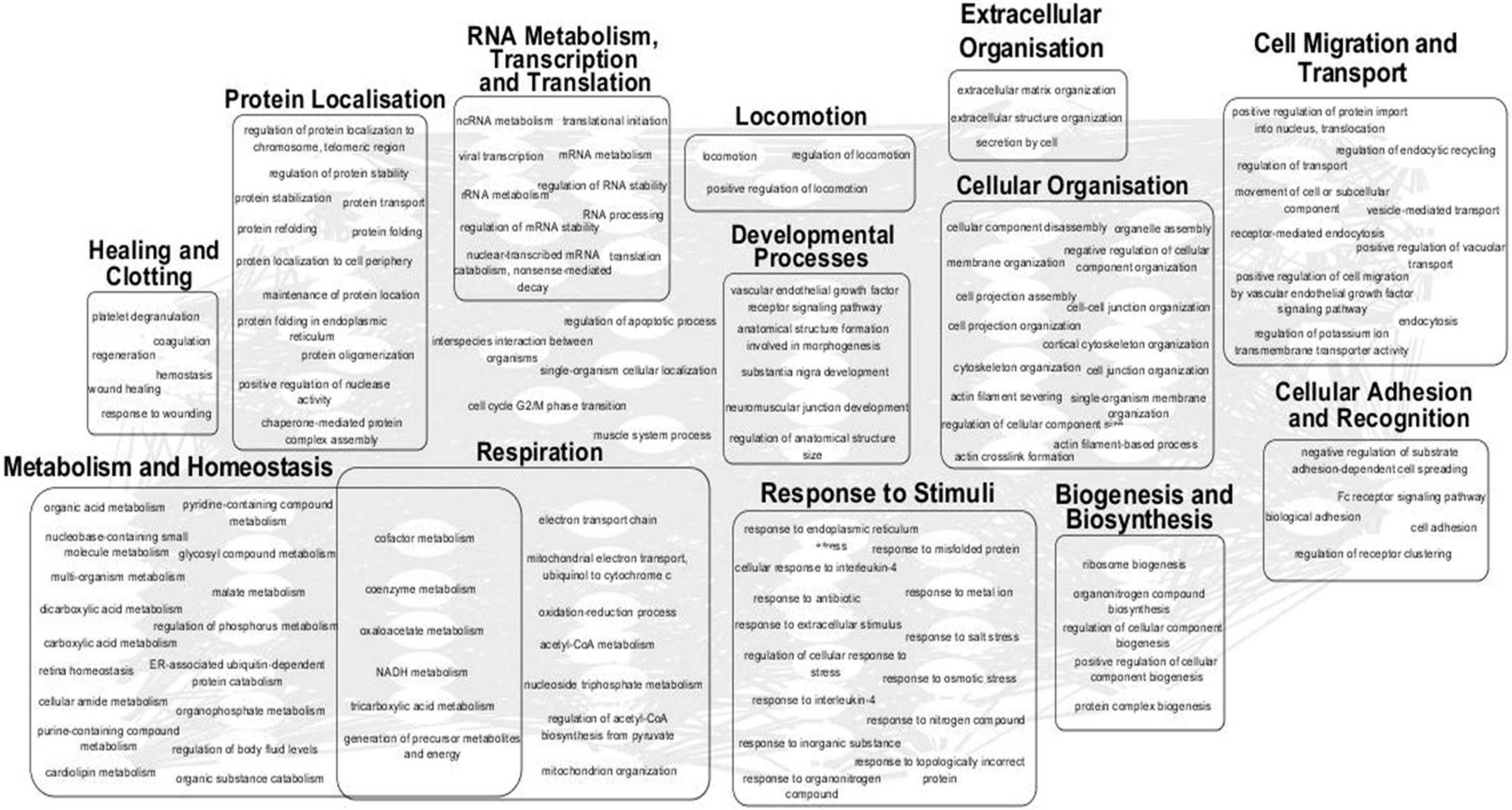


Supplementary Figure 3. Gene Ontology (GO) of upregulated proteins ACL exercise in comparison to ACL control group. ToppGene was used to perform functional enrichment analysis on upregulated proteins to highlight biological processes most. GO Terms (Bonferroni FDR < 0.05) were summarized and visualised using REViGO and Cytoscape. Result shows that the most upregulated proteins involve metabolism, homeostasis and respiration.
