## Supplementary Figure 2 for "The effect of exercise on the protein profile of rat knee joint intra- and extra-articular ligaments"

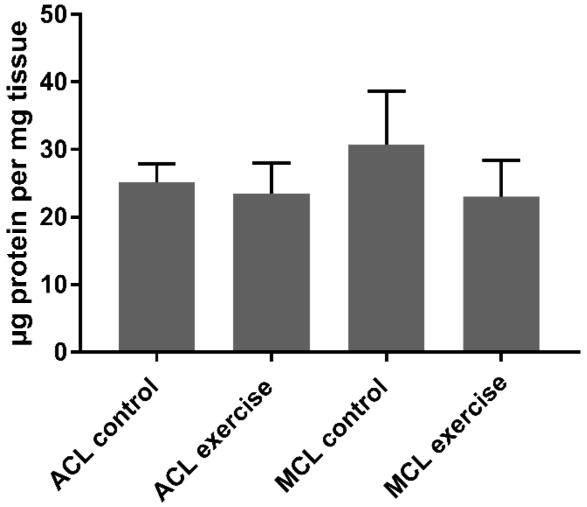


Supplementary Figure 2. (A) Protein concentration yielded between control and exercise groups. Values are mean and error bars represent SD.
