## Supplementary Figure 3 for "The effect of exercise on the protein profile of rat knee joint intra- and extra-articular ligaments"

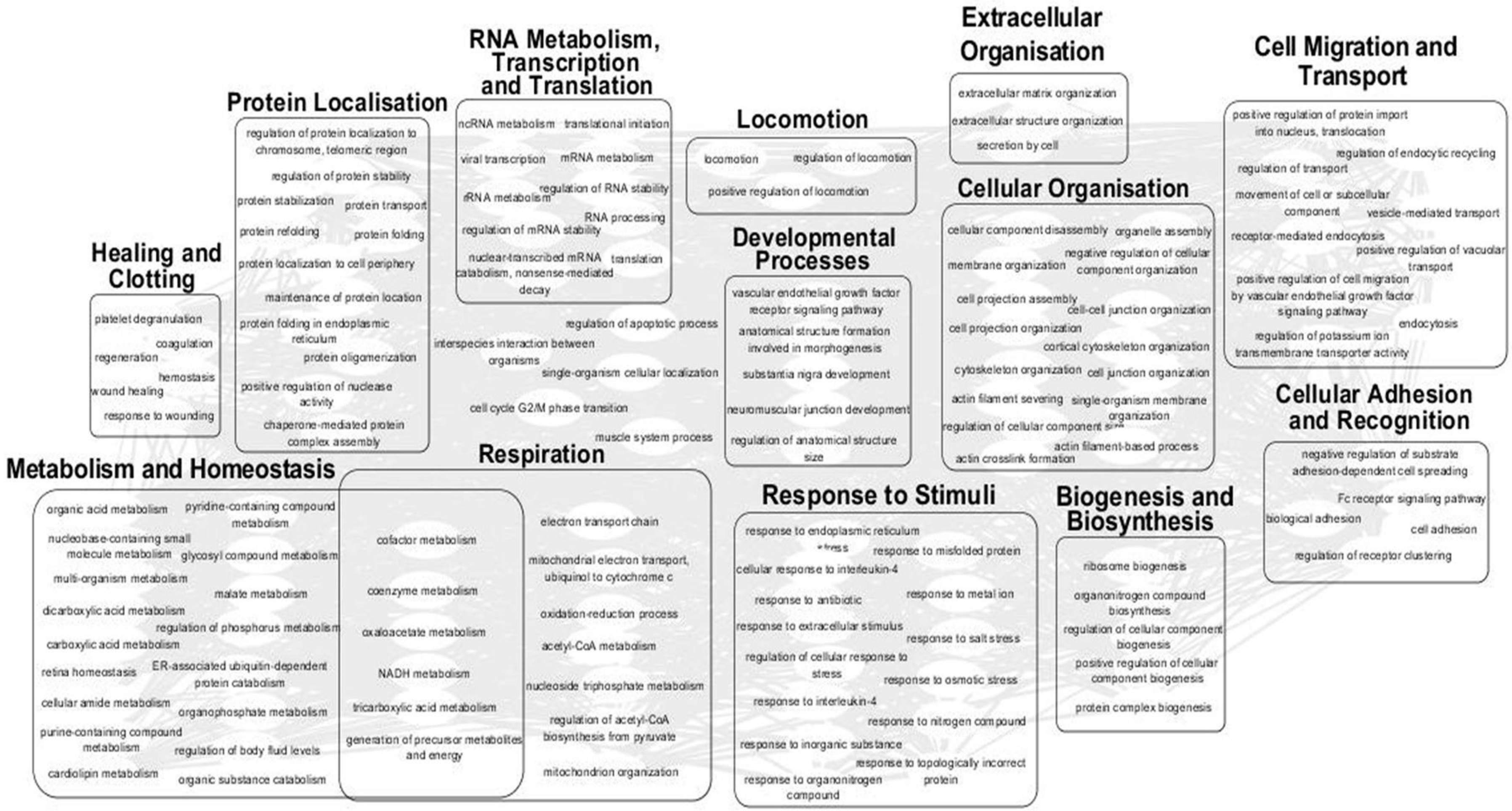


Supplementary Figure 3. Gene Ontology (GO) of upregulated proteins ACL exercise in comparison to ACL control group. ToppGene was used to perform functional enrichment analysis on upregulated proteins to highlight biological processes most. GO Terms (Bonferroni FDR < 0.05) were summarized and visualised using REViGO and Cytoscape. Result shows that the most upregulated proteins involve metabolism, homeostasis and respiration.
